# Rituximab-IgG2 is a phagocytic enhancer in antibody-based immunotherapy of B-cell lymphoma by altering CD47 expression

**DOI:** 10.1101/2024.06.18.599534

**Authors:** Oanh T.P. Nguyen, Sandra Lara, Giovanni Ferro, Matthias Peipp, Sandra Kleinau

## Abstract

Antibody-dependent phagocytosis (ADP) by monocytes and macrophages contributes significantly to the efficacy of many therapeutic monoclonal antibodies (mAbs), including anti-CD20 rituximab (RTX) targeting CD20^+^ B-cell non-Hodgkin lymphomas (NHL). However, ADP is constrained by various immune checkpoints, notably the anti-phagocytic CD47 molecule, necessitating strategies to overcome this resistance.

The IgG2 isotype of RTX induces CD20-mediated apoptosis in B-cell lymphoma cells, and significantly enhances Fc receptor-mediated phagocytosis when used in combination with RTX-IgG1 or RTX-IgG3 mAbs, as previously described. Here, we report that the apoptotic effect of RTX-IgG2 on lymphoma cells contributes to changes in the tumor cell’s CD47 profile by reducing its overall expression and altering its surface distribution. Furthermore, when RTX-IgG2 is combined with other lymphoma-targeting mAbs, such as anti-PD-L1 or anti-CD59, it significantly enhances the ADP of lymphoma cells compared to single mAb treatment.

In summary, RTX-IgG2 acts as a potent phagocytic enhancer by promoting Fc-receptor mediated phagocytosis through apoptosis and reduction of CD47 in malignant CD20^+^ B-cells. RTX-IgG2 represents a valuable therapeutic component in enhancing the effectiveness of different mAbs targeting B-cell NHL.

## Introduction

Among various hematological malignancies classified under non-Hodgkin lymphoma (NHL), B-cell lymphoma is the most prevalent subtype.^1^ In the past decades, the survival rate of B-cell lymphoma patients has been significantly improved thanks to advancements with therapeutic monoclonal antibodies (mAbs). Nevertheless, predicting the treatment responses of most B-cell lymphomas remains challenging due to their significant heterogeneity, highlighting the critical need for ongoing research to improve treatment efficacy and address these complexities.^2^

Since its initial approval in 1997, the chimeric CD20-targeting IgG1 mAb rituximab (RTX) has remained a cornerstone in cancer immunotherapy.^3^ Despite its widespread use, the precise *in vivo* mechanisms of RTX’s action remain uncertain.^4^ Preclinical and clinical data have associated RTX’s efficacy in eliminating CD20^+^ lymphoma cells with various immune responses such as antibody-dependent phagocytosis (ADP), complement-mediated cytotoxicity (CDC), and to a lesser extent, apoptosis.^5–7^ Increasing evidence now suggests that the efficacy of RTX monotherapy is primarily mediated through Fc receptor (FcR)-dependent phagocytosis by monocytes and macrophages.^6,8–13^ However, opsonization with RTX alone often fails to induce a complete eradication of the target cells, indicating the presence of resistance mechanism.^3^

In recent years, a number of important insights into the regulation of phagocytosis have been established. CD47, a transmembrane protein, has been identified as a crucial regulator of phagocytosis by providing an anti-phagocytic (or “don’t-eat-me”) signal. CD47 achieves its anti-phagocytic effect by binding to its cognate receptor – the signal regulatory protein-α (SIRP-α) – which is highly expressed on macrophages.^14,15^ CD47 is ubiquitously expressed in the body and acts as a “marker-of-self”, protecting healthy cells from being phagocytosed, thus maintaining immune homeostasis and preventing autoimmunity.^14^ Senescent or damaged cells, for example erythrocytes, and apoptotic cells have been shown to have reduced CD47 expression or express CD47 with altered conformation. These alterations consequently render the anti-phagocytic CD47/SIRP-α signal ineffective and allow for removal of these cells.^16–19^ Interestingly, recent studies have shown that various malignancies overexpress CD47 as a part of their immune evasion, dampening ADP induced by therapeutic mAbs.^14,15^ This suggests that disrupting CD47-SIRP-α interaction may be pro-phagocytotic, thus potentially improving the therapeutic efficacy of anti-tumor mAbs. Indeed, Métayer *et al.* showed that blocking CD47 greatly increased phagocytosis of B-cell lymphoma cells by macrophages.^20^ Along the same lines, Chao *et al*. reported a synergistic ADP-mediated anti-tumor activity when combining a CD47-blocking mAb with RTX.^13^ Despite these promising findings, clinical trials have revealed significant toxicity associated with CD47-blocking mAbs, including anemia, neutropenia, and thrombocytopenia.^14^ ^21^ Nonetheless, CD47 remains a potential therapeutic target for improving the efficacy of anti-tumor Abs, provided that the therapy of choice selectively targets only CD47 on the target tumor cells.

We have previously demonstrated that the IgG2 isotype of RTX (RTX-IgG2) is capable of inducing CD20-mediated apoptosis in B-cell lymphoma cells.^9^ When combining RTX-IgG2 with the phagocytosis-efficient RTX-IgG1 or RTX-IgG3, it can significantly enhance phagocytosis of B-cell lymphoma cells.^9^ This suggests that RTX-IgG2 presents a promising approach to enhance the efficacy of RTX without contributing to adverse effects.

Here, we aim at investigating the mechanisms underlying the RTX-IgG2-mediated enhancing effect on phagocytosis when combined with RTX isotypes, and further explore its potential when combined with mAbs targeting other lymphoma antigens (PD-L1 and CD59). Our findings reveal, for the first time, that RTX-IgG2-mediated apoptosis is associated with a reduced and altered CD47 expression on CD20^+^ B-cell lymphoma cells, enhancing FcγR-mediated phagocytosis by tumor-specific mAbs.

## Results

### ADP induced by RTX-IgG1 or RTX-IgG3 is enhanced if combined with RTX-IgG2

The capacity of different anti-CD20 RTX isotype variants to stimulate human monocytes to phagocytose CD20^+^ B-cell lymphoma cells varies significantly, and use of certain isotype combinations can further improve the ADP function.^9,10^ Indeed, *in vitro* ADP analyses of human monocytic MonoMac-6 cells, in co-cultures with human CD20^+^ B-cell lymphoma Granta-519 cells (Supplementary Fig. S1) opsonized with different RTX IgG isotypes, demonstrate that RTX-IgG3 is more effective than RTX-IgG1 in inducing phagocytosis (Fig. 1a-b) (Supplementary Fig. S2, S3), regardless of concentration used (Supplementary Fig. S4). In contrast, RTX-IgG2 exhibits quite modest phagocytic activity (Fig. 1a) (Supplementary Fig. S2b). However, when RTX-IgG2 is combined with either RTX-IgG1 or RTX-IgG3, it significantly enhances the ADP response compared with RTX-IgG1 or RTX-IgG3 alone (Fig. 1). RTX-IgG1 achieves an approximately 19% increase in ADP when combined with RTX-IgG2, while RTX-IgG3 shows an increase of about 12% when combined with RTX-IgG2 (Table 1). This enhancing effect of RTX-IgG2 has been associated with CD20-mediated apoptosis, but the specific pathways involved remain unclear.^9^ To address this gap, we have delved deeper into the phagocytic enhancing effect by RTX-IgG2.

**Fig. 1.**
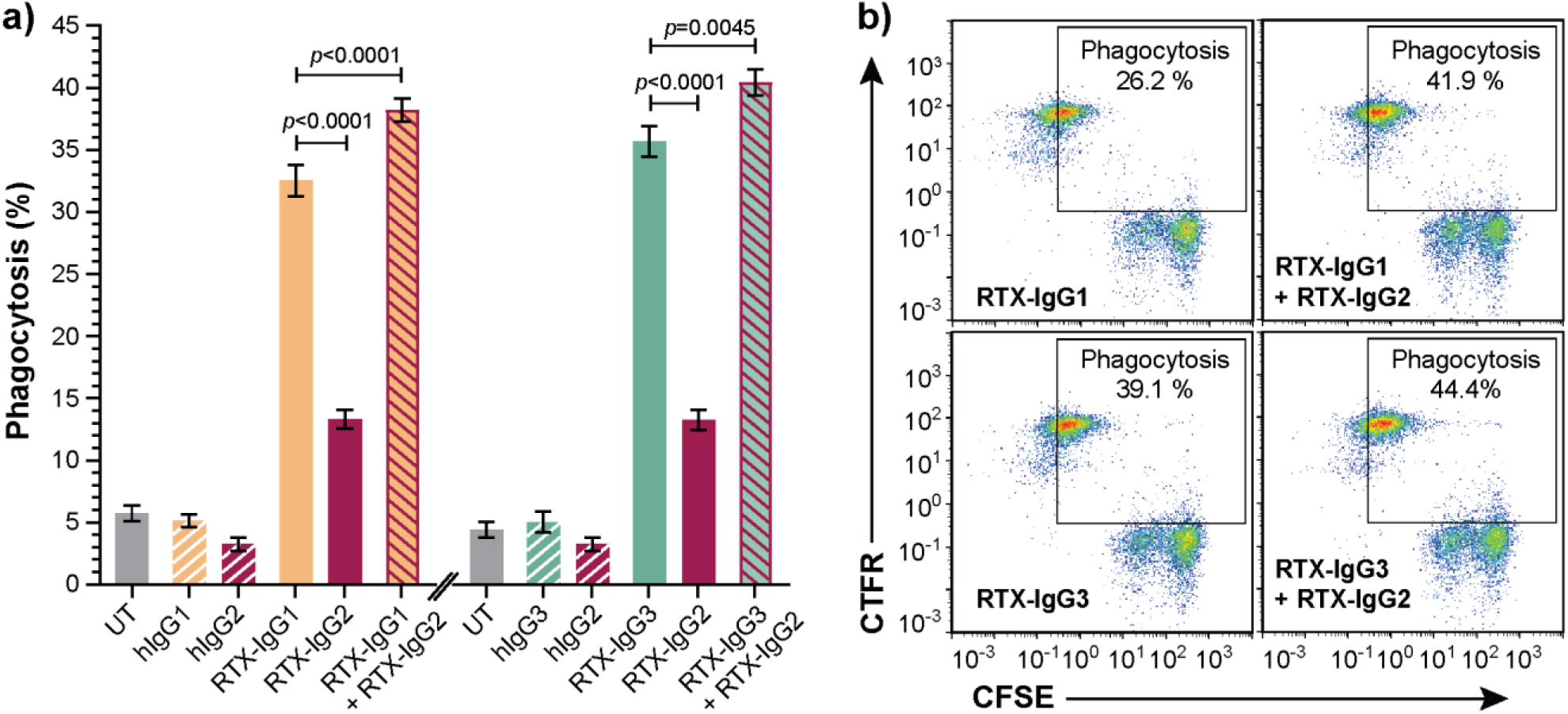
RTX-IgG2 enhances phagocytosis of RTX-IgG1 or RTX-IgG3 treated CD20^+^ B-cell lymphoma cells. Flow cytometry analysis of ADP of Granta-519 B-cell lymphoma cells, treated with anti-CD20 RTX isotypes or human isotype control Abs (1.5 µg/ml), by MonoMac-6 cells at an E:T ratio of 1:1. a) Percentage phagocytosis of Granta-519 cells induced by single RTX-isotypes, or dual combinations of RTX-IgG2 with RTX-IgG1 or RTX-IgG3. Untreated cells (UT), and human Ab isotypes: hIgG1, hIgG2, hIgG3 were used as controls. For dual treatments, Granta-519 cells were pre-opsonized with 1.5 μg/mL of RTX-IgG2 for 30 min followed by 1.5 μg/mL of RTX-IgG1 or RTX-IgG3. Results are shown as mean ± standard error of the mean (SEM) of three independent experiments, each with three biological replicates. Statistical analysis by one-way ANOVA with Tukey-Kramer post-hoc test. b) Representative bivariate plots showing phagocytosis of RTX-treated CFSE-labelled Granta-519 target cells by CTFR-labelled MonoMac-6 effector cells. Phagocytosis was quantified as the percentage of double positive CFSE^+^ CTFR^+^ MonoMac-6 cells (square gate). Increased phagocytosis was observed when RTX-IgG2 was combined with RTX-IgG1 or RTX-IgG3.

**Table 1.** Summary of percentage increase in phagocytosis of CD20^+^ B-cell lymphoma cells (Granta-519) induced by tumor-specific mAb (anti-CD20 RTX, anti-PD-L1, or anti-CD59) in combination with RTX-IgG2 or anti-CD47 mAb (where applicable) in comparison with single use of tumor-specific mAb.

| Treatment | RTX-IgG1 | RTX-IgG3 | αPD-L1 | αCD59 |
| --- | --- | --- | --- | --- |
| +RTX-IgG2 | 19%* | 12% | 31% | 44% |
| +αCD47-fuFc | 52% | 28% | nd | nd |
| +αCD47-siFc | 42% | 17% | nd | nd |

\*Mean (%); fuFc = functional Fc domain; siFc = silenced Fc domain; nd = not determined

### RTX-IgG2 induces significant CD20-mediated apoptosis but minor necrosis

To study the role of apoptosis in RTX-IgG2 mediated enhancement of ADP, we first optimized the concentration of RTX-IgG2 in relation to the apoptosis-inducing agent staurosporine (STR) in Granta-519 cells, while minimizing necrosis. We identified that RTX-IgG2, at concentrations of 1.5-2.5 µg/mL, and STR at 7.5 µM, induced comparable levels of apoptosis (27% ± 1.7 and 26% ± 1.3, respectively), which were significantly higher than apoptosis levels observed in untreated Granta-519 cells (10% ± 1) (Fig. 2a-b) (Supplementary Fig. S5a, S5d). In contrast, RTX-IgG1 and RTX-IgG3 did not induce apoptosis in the Granta-519 cells (Fig. 2a-b). Notably, RTX-IgG2 was able to induce a comparable level of apoptosis as STR, but within a significantly shorter timeframe – 30 min for RTX-IgG2 versus 6 h for STR. Moreover, RTX-IgG2-induced apoptosis was accompanied by negligible levels of necrosis compared to untreated controls (Fig. 2c) (Supplementary Fig. S5b). Similarly, neither RTX-IgG1 nor RTX-IgG3 induced necrosis. STR induced approximately 20% more necrosis in Granta-519 cells compared to untreated controls (Fig. 2c) (Supplementary Fig. S5e), yet the level of necrosis in STR-treated cells remained relatively low, accounting for less than 12% of the total analyzed cell population.

**Fig. 2.**
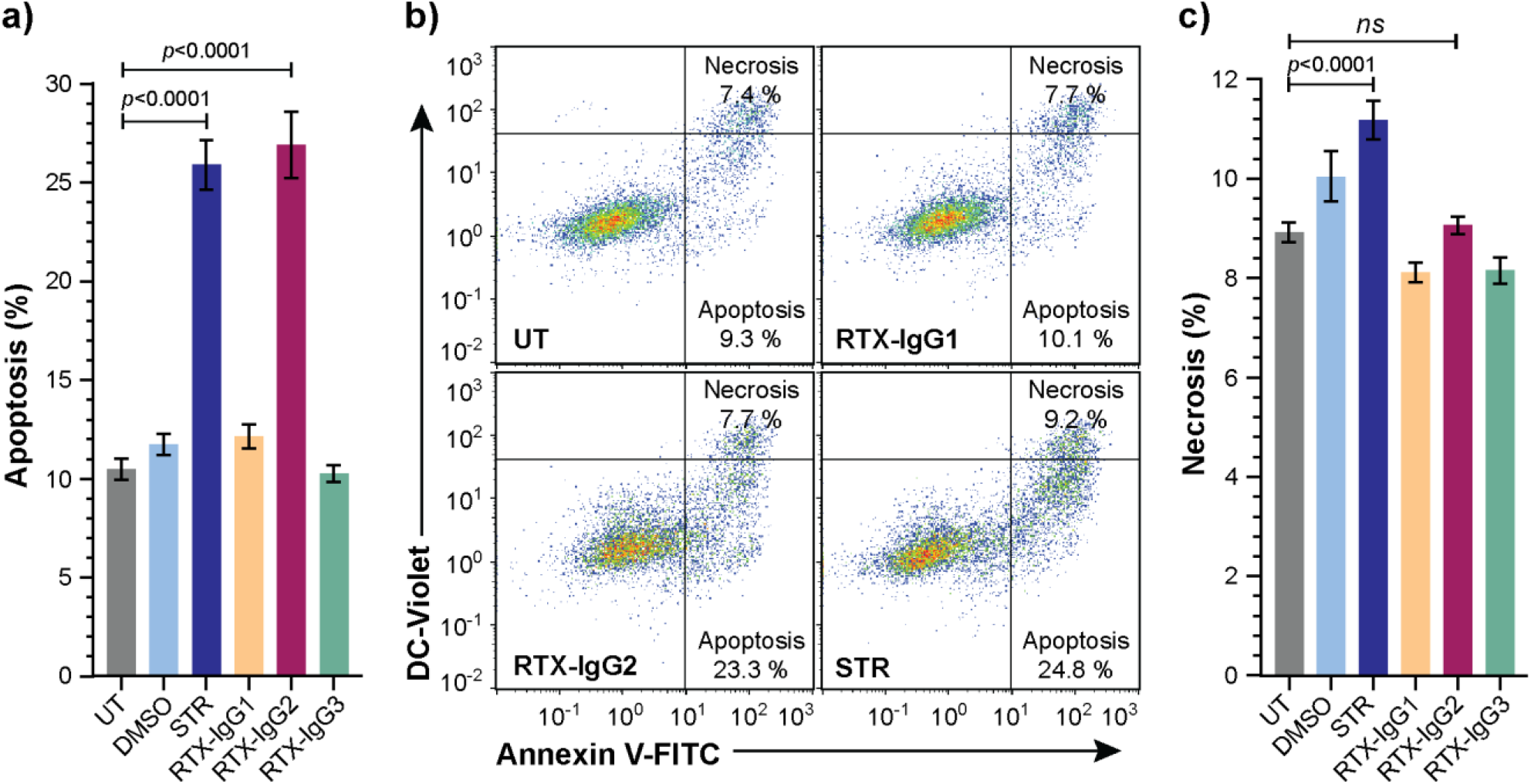
Analysis of cell death in CD20^+^ B-cell lymphoma cells treated with STR or RTX isotypes. Granta-519 cells treated with STR (6 h) or RTX-IgG isotypes (30 min) were analyzed for apoptosis or necrosis compared to untreated cells (UT). Dimethyl sulfoxide (DMSO) was used as vehicle control of STR treatment. a) Percentage apoptosis in UT or treated Granta-519 cells. b) Representative bivariate plots of Granta-519 cells, showing apoptosis and necrosis in UT, and after treatment with RTX-isotypes or STR. Apoptotic cells were identified as Annexin V^+^ DC-Violet^‒^ cells, while double positive (Annexin V^+^ DC-Violet^+^) cells were identified as necrotic cells with compromised cell membrane. c) Percentage necrosis in UT or treated Granta-519 cells. Results in a) and c) show mean ± SEM of three independent experiments, each with three biological replicates. Statistical analysis by one-way ANOVA with Tukey-Kramer post-hoc test (ns = not significant).

Control experiments with a human CD20^‒^ B-cell precursor leukemia cell line (Reh)^22^ (Supplementary Fig. S6a) further affirmed that the apoptosis induced by RTX-IgG2 was CD20-dependent as RTX-IgG2 did not induce apoptosis or necrosis in this CD20^‒^ cell line (Supplementary Fig. S6b-c).

### Apoptosis enhances FcR-mediated phagocytosis

To verify that apoptosis contributes to enhanced ADP, we also investigated the effect of CD20-independent apoptosis, induced by STR, on ADP mediated by RTX isotypes. Indeed, when combined with RTX-IgG1 or RTX-IgG3 treatment on Granta-519 cells, STR significantly enhanced ADP compared to single RTX treatment (Fig. 3). Unaccompanied STR resulted in higher ADP of Granta-519 cells compared to untreated controls, although at inferior levels compared to anti-CD20 mAb single treatment (Fig. 3) (Supplementary Fig. S7).

**Fig. 3.**
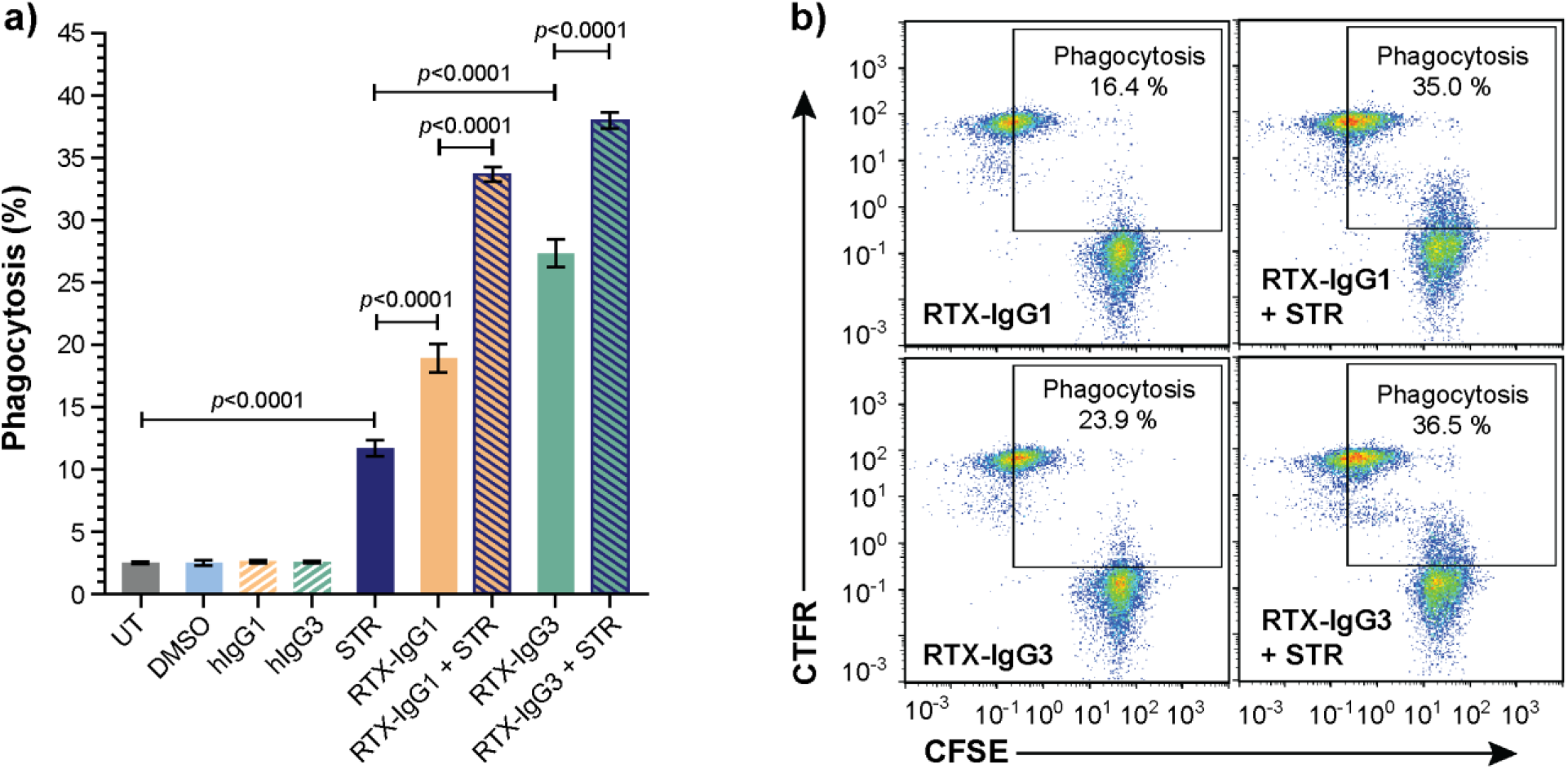
Apoptosis induced by STR in CD20^+^ B-cell lymphoma cells enhances ADP. Flow cytometry analysis of phagocytosis of Granta-519 cells, untreated (UT) or incubated with STR for 6 h before addition of RTX isotypes or isotype controls (1.5 μg/mL), by MonoMac-6 cells (E:T ratio = 1:1). a) Percentage phagocytosis of UT Granta-519 cells, treated with STR, RTX-IgG1 or RTX-IgG3 or combinations thereof. Results shows mean ± SEM of three independent experiments, each with three biological replicates. Statistical analysis by one-way ANOVA with Tukey-Kramer post-hoc test. b) Representative bivariate plots showing phagocytosis of CFSE-labelled Granta-519 target cells, treated with RTX isotypes alone or in combination with STR. Phagocytosis was quantified as the percentage of double positive CFSE^+^ CTFR-MonoMac-6 cells (square gate). Increased phagocytosis was observed when RTX-IgG1 or RTX-IgG3 is combined with STR.

### Apoptosis enhances phagocytosis by impairing “don’t-eat-me” CD47 expression

The level of “don’t-eat-me” antiphagocytic CD47 protein varies on the surfaces of different B-cell lymphoma cell lines (Supplementary Fig. S8a). Among the tested cell lines, Granta-519 cells expressed the highest level of CD47, implying its greater resistance to ADP (Supplementary Fig. S8b). Interestingly, we discovered that apoptosis, induced by either RTX-IgG2 or STR, led to a reduction of CD47 expression in Granta-519 cells (Fig. 4a). STR – being a very efficient apoptosis inducer – significantly reduced the expression level of CD47 to 0.55-fold of the level detected in untreated cells (Fig. 4a-b). Similarly, CD20-mediated apoptosis induced by RTX-IgG2 reduced the expression level of CD47 to 0.78-fold of the level detected in untreated cells, corresponding to a 22% reduction in CD47 on the cell surface (Fig. 4b). The expression pattern of CD47 on the surface of Granta-519 cells was further examined using confocal microscopy. As shown in Fig. 4c (upper panel), CD47 was evenly distributed on the cell membrane of untreated and RTX-IgG1-treated Granta-519 cells. In contrast, RTX-IgG2-or STR-treated Granta-519 cells showed decreased CD47 expression, with CD47 redistributed into scattered clusters on the cell membrane (Fig. 4c; lower panel). Remarkably, while STR and RTX-IgG2 induced similar levels of apoptosis, STR-treated cells exhibited signs of a later stage of apoptosis, such as nuclear deformation and condensed chromatin (visualized by Hoechst 33342 dye), which were absent in RTX-IgG2-treated cells (Fig. 4c, lower panel).

**Fig. 4.**
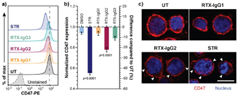
Decreased CD47 expression in RTX-IgG2 treated CD20^+^ B-cell lymphoma cells. CD47 expression was evaluated in Granta-519 cells after incubation with STR (6 h) or treatment with RTX isotypes (1.5 μg/mL) (30 min). a) Representative histograms of CD47 expression in Granta-519 cells after treatment with RTX-IgG1, RTX-IgG2, RTX-IgG3, or STR. Grey dashed line indicates the level of CD47 on untreated cells (UT). b) Bar graph representation of fold change (left Y axis) and percentage difference (right Y axis) in CD47 expression on Granta-519 cells induced by STR or RTX isotypes, normalized to CD47 expression on UT. To obtain the fold change values, mean fluorescence intensity (MFI) of treated samples was first adjusted by subtracting MFI of isotype controls and then normalized to the MFI of UT samples. The percentage difference was calculated by the following formula: ((MFI_treated sample_ – MFI_UT_)*100)/MFI_UT_). Results show mean fold change ± SEM of three independent experiments, each with three biological replicates. Statistical analysis by one-way ANOVA with Tukey-Kramer post-hoc test. c) Confocal microscopy images of CD47 expression in untreated, RTX-IgG1, RTX IgG2 and STR-treated Granta-519 cells. Cells were counterstained with Hoescht 33342 nucleus stain. Disruptions in CD47 cell surface pattern are indicated by white arrows. Scale bar: 10 μm.

### Blocking of CD47 enhances phagocytosis

Our data suggested that decreased levels of CD47 on the surface of Granta-519 cells, induced by RTX-IgG2 or STR, were associated with enhanced phagocytic activity by the MonoMac-6 effector cells. Therefore, in the next experiment, we used CD47-blocking mAbs, either with a functional or a silenced Fc domain, together with RTX-IgG1 or RTX-IgG3. As shown in Fig. 5, blocking CD47 on the Granta-519 target cells significantly enhanced the phagocytic activity by MonoMac-6 cells. When combined with RTX-IgG1 or RTX-IgG3, the CD47-blocking mAb with a functional Fc domain (αCD47-fuFc) increased ADP by 52% and 28%, respectively (Table 1). Meanwhile, the CD47-blocking mAb with a silenced Fc domain (αCD47-siFc) showed a modestly lower, yet comparable, enhancing effect on phagocytosis compared to the αCD47-fuFc, achieving an increase of 42% and 17% when combined with RTX-IgG1 or RTX-IgG3, respectively (Table 1). Notably, both CD47-blocking mAbs could trigger enhanced ADP by themselves in comparison with untreated cells, but at much lower levels than when combined with RTX-IgG1 or RTX-IgG3 (Fig. 5). The αCD47-fuFc mAb alone induced significant phagocytosis (20% ± 3), which was doubled compared to the ADP level induced by αCD47-siFc alone (10% ± 1), underscoring the FcR-mediated phagocytosis induced by the Fc domain of the αCD47-fuFc.

**Fig. 5.**
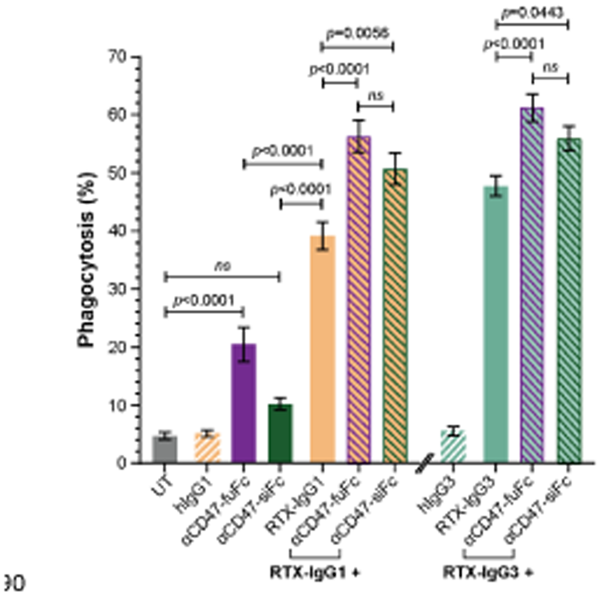
Blocking of CD47 on CD20^+^ B-cell lymphoma cells enhances FcR-mediated ADP. Percentage phagocytosis of Granta-519 cells treated with RTX-IgG1 or RTX-IgG3, and in combination with blocking of CD47 by mouse anti-human CD47 Ab with a functional Fc domain (αCD47-fuFc, 1 μg/mL) or humanized anti-CD47 Ab with a silenced Fc domain (αCD47-siFc, 10 μg/mL), by MonoMac-6 cells (E:T ratio = 2:1). Data are presented as mean ± SEM of three independent experiments, each with three biological replicates. Statistical analysis by one-way ANOVA with Tukey-Kramer post-hoc test (ns = not significant).

### RTX-IgG2 enhances the efficacy of other tumor-targeting mAbs in inducing ADP of B-cell lymphoma cells

Since RTX-IgG2 was capable of altering and reducing the expression of CD47 on the target Granta-519 cells (Fig. 4), we hypothesized that a pre-treatment of Granta-519 cells with RTX-IgG2 would enhance the efficacy of other tumor-targeting mAbs to induce ADP. For this experiment, we chose two target molecules with different expression levels on Granta-519 cells – the immune checkpoint PD-L1 and the complement inhibitor CD59. As shown by Fig. 6a, Granta-519 cells expressed the CD59 antigen markedly, while PD-L1 was weakly expressed (Fig. 6b). When Granta-519 cells were treated with RTX-IgG2 in combination with mAbs targeting PD-L1 (αPD-L1) or CD59 (αCD59) antigens, the level of phagocytosis increased significantly compared to cells treated with αPD-L1 or αCD59 mAb alone (Fig. 6c-d). Notably, the mAb targeting the highly expressed CD59 antigen mediated a significant phagocytosis alone, in contrast to the mAb reactive to the low-expressing PD-L1 molecule. Correspondingly, RTX-IgG2 enhanced the phagocytic efficacy of αCD59 by 44% and of αPD-L1 by 31% (Table 1).

**Fig. 6.**
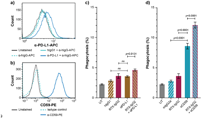
RTX-IgG2 enhances phagocytosis of CD20^+^ B-cell lymphoma cells induced by anti-PDL1 or anti-CD59 mAb. Representative histograms of a) PD-L1 and b) CD59 expression on Granta-519 cells. c-d) Percentage phagocytosis of Granta-519 cells by MonoMac-6 cells (E:T ratio = 1:1), induced by c) human anti-PDL1 mAb (αPD-L1, 1.5 μg/mL) or d) mouse anti-human CD59 mAb (αCD59, 2.5 μg/mL) alone, or in combination with a pre-treatment with RTX-IgG2. Data are presented as mean ± SEM of three independent experiments, each with three biological replicates. Statistical analysis by one-way ANOVA with Tukey-Kramer post-hoc test (ns = not significant).

## Discussion

Macrophages are the most abundant tumor-infiltrating immune cells in most human solid tumors, making them appealing therapeutic targets for cancer therapy.^23,24^ However, their role as effector cells in cancer therapy remains underappreciated due to their intricate and polarized roles within the tumor microenvironment. A number of studies have associated a high content of tumor-infiltrating macrophages with unfavorable prognoses, especially in patients treated with certain chemotherapeutic agents.^25–27^ Conversely, macrophages have been shown to contribute significantly to the efficacy of mAb-based cancer immunotherapies through ADP.^8,9,28–31^ In fact, macrophage infiltration has been shown to improve therapeutic responses in patients receiving a combined regimen of tumor-targeting Ab and chemotherapy.^32^ Additionally, monocytes – which express activating FcRs such as FcγRI (CD64) and FcγRIIa (CD32a) – have demonstrated the ability to eliminate tumor cells through ADP as well.^9–11^ These lines of research necessitate further investigation on monocytes/macrophages-based cancer therapies to unlock their full anti-tumor potential.

Phagocytosis by macrophages/monocytes is mainly govern by the ubiquitously expressed “don’t-eat-me” CD47. Overexpression of CD47 has been shown to associate with adverse prognosis in mantle cell lymphoma patients and play an important role in dissemination of B-cell NHL. ^13^ ^33^ Strategies targeting the interaction between CD47 and its receptor SIRP-α have demonstrated promising results either as monotherapies or in combination with other tumor-targeting therapies.^34^ However, anti-CD47 therapies still encounter many setbacks in terms of selectivity, efficacy, and safety profile as CD47 is expressed not only on tumor cells but also on non-malignant cells.

In this study, utilizing the CD20^+^CD47^high^ Granta-519 lymphoma B-cell model, we found that CD20-mediated apoptosis induced by RTX-IgG2 resulted in a reduction of CD47 expression on the Granta-519 cells. We also demonstrate that RTX-IgG2-treated lymphoma cells become significantly more susceptible to RTX-IgG1- or RTX-IgG3-mediated phagocytosis, which is consistent with our previous results.^9^ The apoptosis-inducer STR generated comparable levels of apoptosis as RTX-IgG2 and, similarly to RTX-IgG2, caused a reduction in CD47 expression, confirming the cause-effect relationship between apoptosis and CD47 reduction. STR also exhibited equivalent enhancing effect on phagocytosis when combined with RTX-IgG1 or RTX-IgG3. In support of our data, similar apoptosis associated decreased CD47 expression has been reported in various cell types by Gardai *et al.*^16^ It is important to note that STR-induced effects are non-selective and independent of CD20, resulting in high levels of background apoptosis. In contrast, RTX-IgG2 specifically targets only CD20^+^ cells.

While apoptosis induction by RTX-IgG2 has previously been reported, its exact mechanism remains incompletely understood.^9,35^ Compared to RTX-IgG1, RTX-IgG2 induces similar level of homotypic adhesion (Supplementary Fig. S3) (Supplementary Video S1), while exhibiting a substantially reduced ability to induce ADP and CDC.^6,9,35^ In fact, RTX-IgG2 binds CD20^+^ B-cells at only half the density of RTX-IgG1 and RTX-IgG3 (Supplementary Fig. S9),^35^ but induces significantly more apoptosis in CD20^+^ B-cell lymphoma cells.^9,35^ For these reasons, we propose that RTX-IgG2, although binding to the same CD20 epitope as RTX-IgG1, attaches in different binding modes and geometries, in support of the hypothesis by Konitzer *et al*.^35^ Among all IgG isotypes, IgG2 possesses the most rigid hinge region and unique alternative covalent links between its Fab domains and the hinge.^36,37^ Given the significant differences in the hinge region of IgG2, it is reasonable to speculate that these distinct structural features may affect its binding geometry to target receptor.^35,38–40^ Indeed, recent studies have correlated the rigidity of the IgG2 hinge with agonistic function,^38,41,42^ suggesting that mAbs of the IgG2 isotype can elicit agonistic activity upon target binding by closely crosslinking target receptors, thereby promoting downstream signaling.^38–43^ Based on these literatures, we attribute the apoptosis-inducing ability of RTX-IgG2 to its unique hinge structure and the resulted agonistic activity.

To our knowledge, our experiment is the first that associates RTX-IgG2-mediated apoptosis with a reduced CD47 expression and an enhancing effect on phagocytosis. We also reveal with microscopic analysis that the pattern of CD47 expression on the surface of RTX-IgG2-treated cells shifts from a homogeneous distribution to a more clustered arrangement. A similar change in spatial distribution of CD47 has previously been reported in aged mouse erythrocytes and human Jurkat T-cells.^16,19^ These studies, together with our observation, suggest that apoptosis trigger major structural alterations of the cell plasma membrane, which may either destabilize lateral molecular interactions needed for the proper CD47 “don’t-eat-me” signaling^16,19^ or induce conformation changes of CD47, hindering its interaction with SIRP-α^44^.

Based on these findings, we hypothesized that a pre-treatment with the agonistic RTX-IgG2 – which reduces the anti-phagocytic CD47 on the surface of target cells – may enhance the phagocytic efficacy of other tumor-targeting mAbs. Indeed, we observed significantly increased phagocytosis of Granta-519 cells treated with anti PD-L1 or anti-CD59 mAbs. Notably, a higher level of phagocytosis correlated with the expression level of the target molecule, as illustrated by CD59. In fact, the potency of RTX-IgG2 as a phagocytic enhancer might be even better when it does not need to compete with mAbs targeting the same antigen/epitope (CD20). Collectively, our results support the role of RTX-IgG2 as an exclusive and target-specific enhancer for FcR-mediated phagocytosis of CD20^+^ B-cell lymphoma cells.

Considering that we associated the reduction in CD47 expression on the Granta-519 cells with an enhanced phagocytosis, we conducted a CD47-blocking experiment to further validate our findings. For this experiment, we used two different versions of CD47-blocking mAbs: a commercially available mouse IgG2a anti-human CD47 Ab (αCD47-fuFc) and an engineered humanized IgG2 anti-CD47 Ab harboring a completely silenced Fc domain (αCD47-siFc).^45^ Both CD47-blocking mAbs effectively enhanced the phagocytic uptake of Granta-519 cells when combined with RTX-IgG1 or RTX-IgG3 (Table 1). However, αCD47-fuFc alone exerted notable Fc-mediated phagocytosis, suggesting potential on-target, off-tumor effects. The pronounced phagocytosis induced by αCD47-fuFc was likely mediated by its Fc domain, complicating the assessment of whether a synergistic effect between this Ab and RTX-IgG1 or RTX-IgG3 was achieved. In contrast, the αCD47-siFc alone induced negligible phagocytosis, indicating effective abrogation of undesired Fc-mediated engagement. This experiment conclusively confirmed that blocking CD47 can synergistically enhance RTX-mediated phagocytosis, provided that the CD47-blocking mAb is carefully designed to avoid undesired Fc receptor engagement on effectors cells and consequent toxicity. As agonistic activity of IgG2 has been demonstrated to occur independently of FcγR engagement,^39^ further utilization of αCD47-siFc warrants comprehensive analyses of its potential agonistic effects.

In conclusion, our study demonstrates that RTX-IgG2 enhances ADP of CD20^+^ B-cell lymphoma cells through CD20-mediated apoptosis and reduction of CD47. This finding emphasizes the potential of RTX-IgG2 as a valuable agonist in B-cell NHL treatment. Combining RTX-IgG2 with the clinical standard RTX-IgG1, or potentially any other lymphoma-targeting antibody, offers an exclusive target specific approach to increase the efficacy of CD20^+^ B-cell lymphoma therapies.

Our findings also suggest that RTX-IgG2 could potentially enhance the effectiveness of RTX therapy in treating autoimmune B-cell disorders. Furthermore, strategic development of tumor-specific IgG2 mAbs with apoptotic capacity may offer a promising avenue for advancing antibody-based immunotherapy.

## Methods

### Cell culture

The human acute monocytic leukemia cell line MonoMac-6, authenticated by STR-profiling (Microsynth AG, Balgach, Switzerland), was used as monocytic effector cells in the ADP assay. The MonoMac-6 cells express the phagocytosis-regulating receptor FcγRIIa (CD32a) (Supplementary Fig. S10a) as well as FcγRI (CD64),^9^ and display the signal-regulatory protein α (SIRP-α) (Supplementary Fig. S10b). MonoMac-6 cells were maintained in suspension cell culture flask (cat. no. 3910.502, Sarstedt AC & Co KG, Nümbrecht, Germany) using complete Roswell Park Memorial Institute medium (cRPMI) that contained RPMI 1640 (cat. no. 21875034, Gibco, Thermo Fisher Scientific), 10% heat-inactivated (h.i.) fetal bovine serum (FBS) (cat. no. 11550356, Gibco, Thermo Fisher Scientific), and 1% Penicillin-Streptomycin (PenStrep) (cat.no. P4333, Merck, Darmstadt, Germany).

The human B-cell lymphoma cell line Granta-519, with distinct expression of CD20 (Supplementary Fig. S1), was utilized as tumor target cells. They originate from a mantle cell lymphoma of a 58-year-old female patient in 1991 and was purchased from the German Collection of Microorganisms and Cell Culture (cat. no. ACC 342, DMSZ, Braunschweig, Germany). The Granta-519 cells were cultured in suspension cell culture flask (cat. no. 3910.502) using Dulbecco’s Modified Eagle Medium (DMEM) (cat. no. 11965092, Gibco, Thermo Fisher Scientific, Waltham, MA, USA) with addition of 10% h.i FBS and 1% PenStrep (complete DMEM; cDMEM).

The human B-cell precursor leukemia cell line Reh, lacking the expression of CD20 (Supplementary Fig. S6a), was kindly provided by Ola Söderberg (Department of Pharmaceutical Biosciences, Uppsala University) and maintained in cRPMI.

All cells were cultured in a humidified cell culture incubator (InCuSafe, LabRum, Östersund, Sweden) at 37°C under 5% CO_2_ and were routinely tested for mycoplasma contamination using MycoStrip™ detection kit (cat. no. rep-mys, InvivoGen, San Diego, CA, USA).

### Antibodies

The human anti-CD20 (RTX) isotype family; IgG1, IgG2, IgG3, and IgG4 anti-CD20, was purchased from InvivoGen (cat. no. hcd20-mab1, hcd20-mab2, hcd20-mab3, and hcd20-mab4). Human recombinant isotype controls (kappa allotype); IgG1, IgG2, IgG3, and IgG4 were obtained from BioRad (Hercules, CA, USA) (cat. no. HCA192, HCA193, HCA194, HCA195). Blocking of CD47 was performed with purified mouse anti-human CD47 mAb (αCD47-fuFc) clone CC2C6 (cat. no. 323102, Biolegend, San Diego, CA, USA), and humanized anti-CD47 IgG2σ mAb (αCD47-siFc) – a variant of magrolimab harboring a completely silenced Fc.^45^

For cell surface staining, the following mAbs were used: APC-conjugated mouse IgG2a anti-human CD20 (clone LT20, cat. no. H12155A, EuroBioScience, Friesoythe, Germany), PE-conjugated mouse IgG1 anti-human CD47 (clone CC2C6, cat. no. 323108, Biolegend), PE-conjugated mouse IgG2a anti-human CD59 (clone p282 (H19), cat. no. 304707, Biolegend), recombinant human IgG1 anti-human PD-L1 (cat. no. hpdl1-mab1, InvivoGen), APC-conjugated mouse anti-human IgG secondary Ab (cat. no. 562025, BD Biosciences, Franklin Lakes, NJ, USA), APC-conjugated mouse IgG2a anti-human SIRP-α Ab (clone 15-414, cat. no. 372109, Biolegend), FITC-conjugated mouse IgG2b anti-human CD32 Ab (clone IV.3, cat. no. 60012Fl.1, StemCell Technologies, Vancouver, BC, Canada). Mouse isotype controls APC-conjugated mouse IgG2a, PE-conjugated mouse IgG1, PE-conjugated mouse IgG2a, and FITC-conjugated mouse IgG2b (cat. no. C12386A, C12385B, C12386P, C12387F) were purchased from EuroBioScience.

### Antibody-dependent phagocytosis (ADP)

MonoMac-6 effector cells were stained with 1 μM of CellTrace™ Far Red dye (CTFR, cat. no. C34564, Thermo Fisher Scientific) in PBS (cat. no. M09-9400-100, Medicago, Quebec City, QC, Canada,) at 10^6^ cells/mL for 20 min at 37°C. After wash with pre-warmed cRPMI (to remove excessive dye), the cells were adjusted to 7.5×10^5^ cells/mL and stimulated with recombinant human interferon gamma (IFNγ, cat. no. PHP050, BioRad) at 0.2 μg/mL for 3 h at 37°C to activate the phagocytic ability of MonoMac-6.^10^ Meanwhile, Granta-519 target cells were stained with Vybrant™ CFDA SE dye (CFSE, cat. no. V12883, Thermo Fisher Scientific) following manufacturer’s instruction prior seeding for monolayers at a cell density of 7.5×10^4^ cells/100 μL per well in a round bottomed 96-well plate (cat. no. 83.3925, Sarstedt). After 3 h, Mono-Mac 6 cells were harvested, resuspended at 7.5×10^4^ or 15×10^4^ cells/100 μL per well (for an effector to target (E:T) ratio of 1:1 or 2:1, respectively), and added to the target cells for co-culturing. For the co-cultures, both cell types were resuspended in an assay medium (cDMEM_low-Glu_) which contained low-glucose (1 g/L) DMEM (cat. no. 31885023, Gibco, Thermo Fisher Scientific), 10% h.i. FBS, and 1% PenStrep. Co-cultures of effectors and target cells were immediately incubated with RTX or isotype control Abs at a concentration of 1.5 μg/mL, and kept at 37°C for 1 h. Untreated cells were used as negative controls. The concentration of RTX was selected from prior testing for optimal induction of phagocytosis (Supplementary Fig. S4). After 1 h, the plate was centrifuged at 500 ×g for 3 min at 4°C and the supernatant was removed. Cells were washed twice with 250 μL of cold PBS containing 0.5% h.i. FBS (flow staining buffer). After the last washing step, cells were resuspended in 100-200 μL of cold flow staining buffer, kept on ice, and analyzed immediately using a MACSQuant VYB flow cytometer (Miltenyi Biotec, Bergisch Gladbach, Germany). Before analyzing cell samples, the flow cytometer was calibrated with MACSQuant Calibration Beads (cat. no. 130-093-607, Milteny Biotec) and compensation was performed using unstained and single-stained samples. A minimum of 15’000 events were acquired for each sample. CTFR^+^CFSE^+^ MonoMac-6 cells were considered positive phagocytic cells (gating strategy shown in Supplementary Fig. S11).

For enhanced phagocytosis assays, target cells (7.5×10^4^ cells/100 μL per well) were pre-incubated with RTX-IgG2 (1.5 μg/mL), αCD47-fuFc (1 μg/mL), or αCD47-siFc (10 μg/mL), for 30 min before addition of RTX-IgG1 or RTX-IgG3 and MonoMac-6 cells. In additional experiments, the same amount of target cells per well was treated with staurosporine (STR) (cat. no. S1421, Selleck Chemicals, Houston, TX, USA) at 7.5 μM for 6 h at 37°C prior to stimulation with RTX-IgG1 or RTX-IgG3. Dimethyl sulfoxide (DMSO) (cat. no. D8418, Merck) was used to prepare the STR stock solution and thus, was used as vehicle control for these experiments.

### Apoptosis assay and CD47 expression analysis

Granta-519 cells were seeded in monolayers at a cell density of 7.5×10^4^ cells/100 μL per well in a round bottomed 96-well plate prior to treatment. The seeded cells were incubated with 7.5 μM STR for 6 h or 1.5 μg/mL of RTX isotypes for 30 min at 37°C. After centrifugation for 5 min at 600 ×g at 4°C, cells were washed with cold flow staining buffer, resuspended in 50 μL of flow staining buffer containing 1× LIVE/DEAD™ Fixable Violet dye (DC-Violet) (cat. no. L34963, Thermo Fisher Scientific) and 1× Human TruStain FcX™ (Fc Receptor Blocking Solution, cat. no. 422302, Biolegend), and kept at room temperature (RT) for 10 min. Positive controls for DC-Violet were prepared by treating Granta-519 cells with 90% ethanol (cat. no. 20821.310, VWR, Radnor, PA, USA) for 1 min. Subsequently, PE-IgG1 anti-CD47 and PE-IgG1 isotype control Abs were added and the cells were stained for 30 min on ice. Cells were subsequently centrifuged at 600 ×g at 4°C, washed with flow staining buffer, resuspended in 100 μL of FITC-conjugated Annexin V (cat. no. 640906, Biolegend) prepared in 1× Annexin V binding buffer (cat. no. 556454, BD Biosciences), and stained for 15 min at RT. After adding 100 μL of 1× Annexin V binding buffer per well, cells were collected and analyzed using the MACSQuant VYB flow cytometer. Unstained and single-stained samples were used for compensation and gating. A minimum of 10’000 events were acquired for each sample. DC-Violet^+^ cells were considered necrotic cells and thus, excluded from further analyses. All DC-Violet^−^ cells were analyzed for CD47 expression, and Annexin V^+^ DC-Violet^−^ Granta-519 cells were considered apoptotic cells (gating strategy shown in Supplementary Fig. S12).

### Immunofluorescence staining and fluorescence microscopy

For live-cell imaging, Granta-519 and MonoMac-6 were prepared according to the ADP protocol before loaded into a flat-bottomed 96 well plate (cat. no. 32096, SPL Life Sciences Co., Ltd., Gyeonggi-do, Korea). Live-cell imaging was performed for 1 h on a fluorescence Nikon Eclipse Ti microscope (Nikon Europe B.V., Amsterdam, Netherlands) with 10-min intervals using a Plan Apo 10×/0.45 objective.

To analyze CD47 expression, RTX or STR-treated Granta-519 cells were washed with cold flow staining buffer and fixed with 50 μL of fixation buffer (cat. no. 88-8824, Thermo Fisher Scientific) for 15 min on ice. The fixed cells were washed twice with cold flow staining buffer and blocked by 1% bovine serum albumin (BSA) (Fraction V, cat. 422371X, VWR) in PBS for 1 h. Next, the cells were incubated with 3 μg/mL of CF®640R-conjugated mouse anti-human CD47 Ab (clone B6H12.2, cat. no. BNC400437-100, Biotium, Fremont, CA, USA) and 1× Hoechst 33342 Ready Flow™ Reagent (cat. no. R37165, Thermo Fisher Scientific), prepared in 1% BSA in PBS overnight at 4°C. Stained cells were then washed, centrifuged, re-dispersed in 20 μL of deionized water, and added on to Superfrost plus microscope slides (cat. no. cat. J1800AMNZ, Thermo Fisher Scientific) for air-drying. Glass coverslips (#1.5, cat. no. 12323148, Fisher Scientific, Hampton, NH, USA) were mounted on the microscope slides using Epredia™ Immu-Mount™ mounting medium (cat. no. 10662815, Fisher Scientific). Fluorescence images of mounted cells were acquired with a Zeiss LSM 700 confocal microscope (Carl Zeiss AG, Oberkochen, Germany) using a Plan-Apochromat 63×/1.4 Oil DIC M27 objective.

### Data analysis

Microscope images were processed and analyzed by ZEISS ZEN lite (black edition) software (Carl Zeiss AG) or the open source Java application ImageJ (https://imagej.nih.gov/ij/). Flow cytometry data were analyzed by FlowJo 10.9.0 software (BD Biosciences). Data were statistically analyzed by one-way ANOVA with Tukey-Kramer post-hoc test and visualized using GraphPad Prism 9.0.0 software (GraphPad Software, Boston, MA, USA). All results are displayed as mean ± standard error of the mean (SEM) of three independent experiments, with a significance of p < 0.05 unless indicated differently. All data were visualized by Adobe Illustrator (Adobe Inc., San Jose, CA, USA) using files from aforementioned software.

## Data availability

All data relevant to the study are included in the article or uploaded as supplementary information. Further inquiries can be directed to the corresponding author.

## Acknowledgements

The authors thank the staff at the BioVis facility, especially Jeremy Adler (Rudbeck Laboratory, Uppsala University, Uppsala, Sweden) and Inkyung Jang (Department of Cell and Molecular Biology, Uppsala University) for technical support. This work was financially supported by Uppsala University.

## Contributions

S.K. conceived the study and together with O.T.P.N designed the experiments. S.L. and G.F. performed preliminary experiments. O.T.P.N performed experiments and analyzed data. M.P. produced and contributed with biological material. O.T.P.N and S.K. wrote and revised the manuscript. All authors approved the submitted version of the manuscript.

