## Supplementary information for "Rituximab-IgG2 is a phagocytic enhancer in antibody-based immunotherapy of B-cell lymphoma by altering CD47 expression"

### Supplementary figures

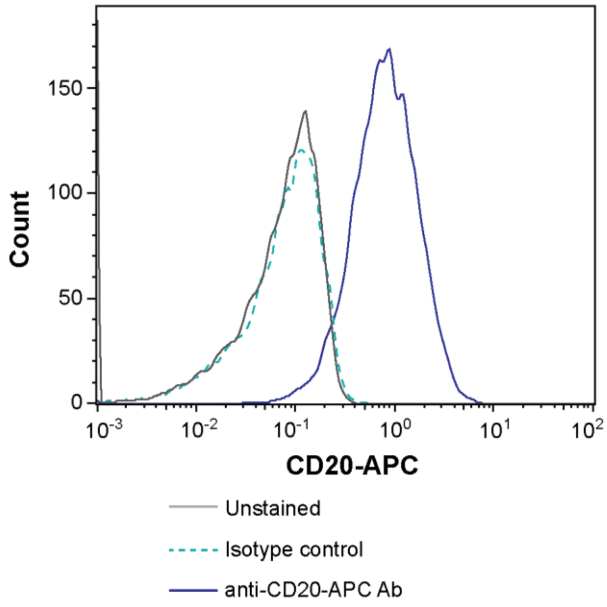


**Supplementary Fig. S1.** Representative histogram of CD20 staining of Granta-519 cells.

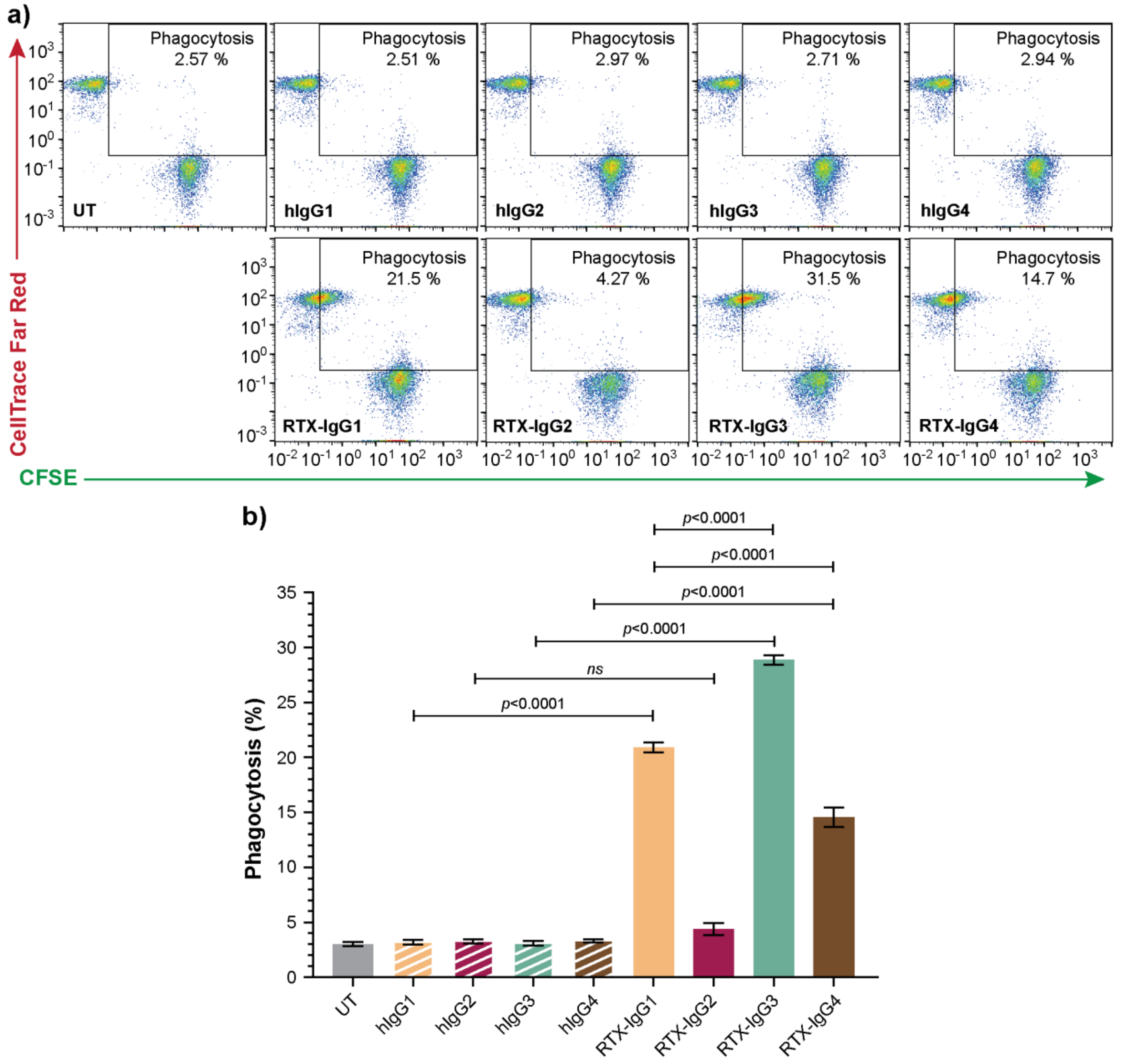


**Supplementary Fig. S2.** a) Representative flow plots showing ADP of Granta-519 cells treated with RTX isotypes (RTX-IgG1-4) or isotype control Abs (hIgG1-4), by MonoMac-6 cells (E:T ratio = 1:1). The phagocytosis of CFSE-labelled Granta-519 cells was quantified as the percentage of double positive $CFSE^{+}CTFR^{+}$ MonoMac-6 cells (rectangular gate). The gating was set based on unstained and single-colored stained controls. b) Percentage phagocytosis of Granta-519 cells, induced by single RTX-isotypes (IgG1-4) or human isotype controls (hIgG1-4), by MonoMac-6 cells (E:T ratio = 1:1). Untreated cells (UT) were used as controls. Results are shown as mean ± SEM of three independent experiments, each with three biological replicates. Statistical analysis by one-way ANOVA with Tukey-Kramer post-hoc test (ns = not significant).

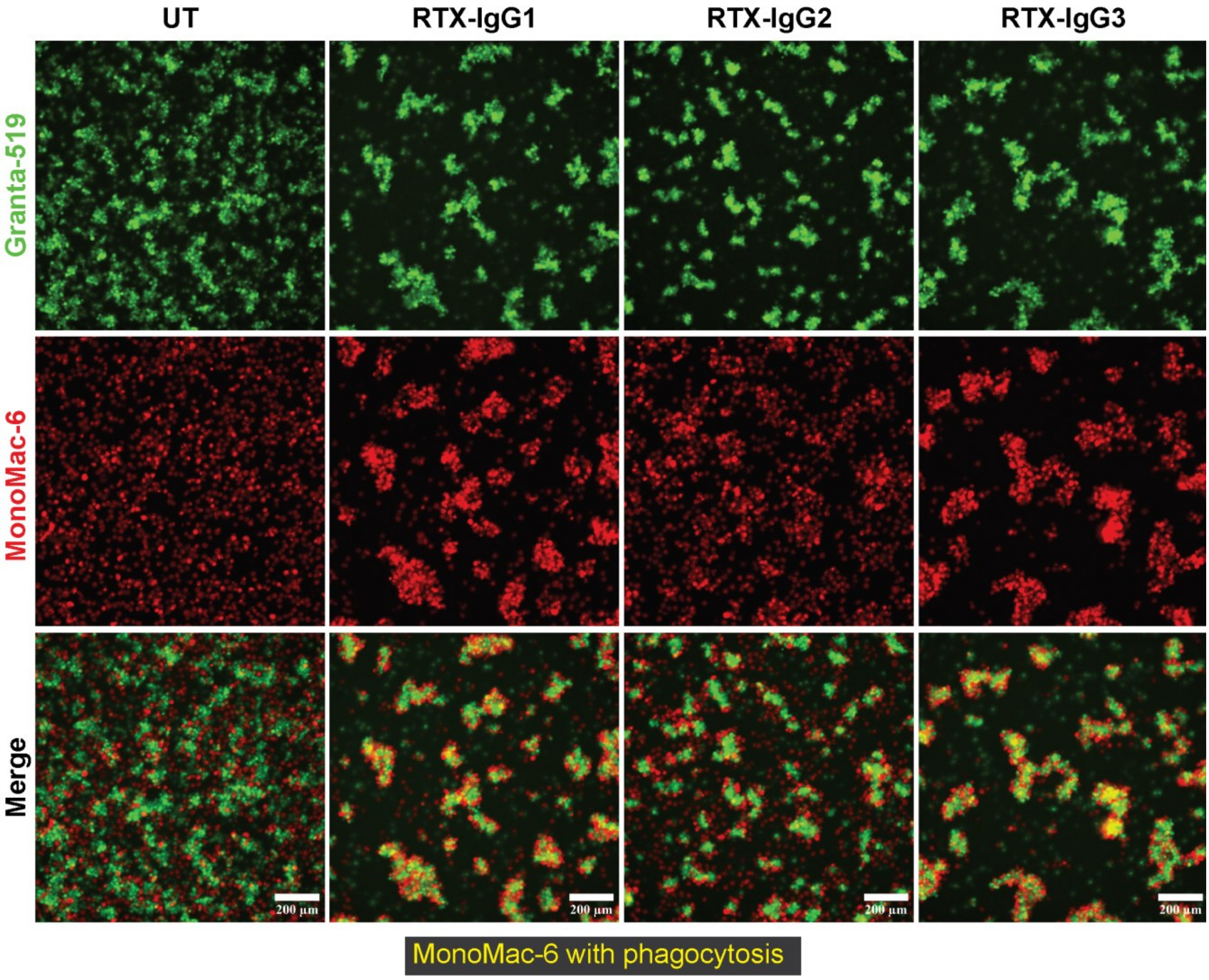


**Supplementary Fig. S3.** Microscopic analysis of ADP of $CD20^+$ B cell lymphoma cells (Granta-519), induced by RTX-IgG1, RTX-IgG2, or RTX-IgG3 by MonoMac-6 cells (E:T ratio = 1:1). All tested RTX isotypes induced a comparable level of homotypic adhesion in CFSE-labeled Granta-519 cells (top panel). The majority of CTFR-labeled MonoMac-6 were attracted to the vicinity or attached to Granta-519 cell clusters in the presence of RTX-IgG1 or RTX-IgG3 (middle panel), with many MonoMac-6 cells becoming double positive for CFSE and CTFR (yellow color) (bottom panel), indicating efficient phagocytosis. In contrast, significantly less MonoMac-6 cells attached to RTX-IgG2-treated Granta-519 cell clusters (middle panel). Only a few $CFSE^+CTFR^+$ MonoMac-6 cells were observed in untreated (UT) or RTX-IgG2-treated co-cultures (bottom panel). Scale bars: 200µm.

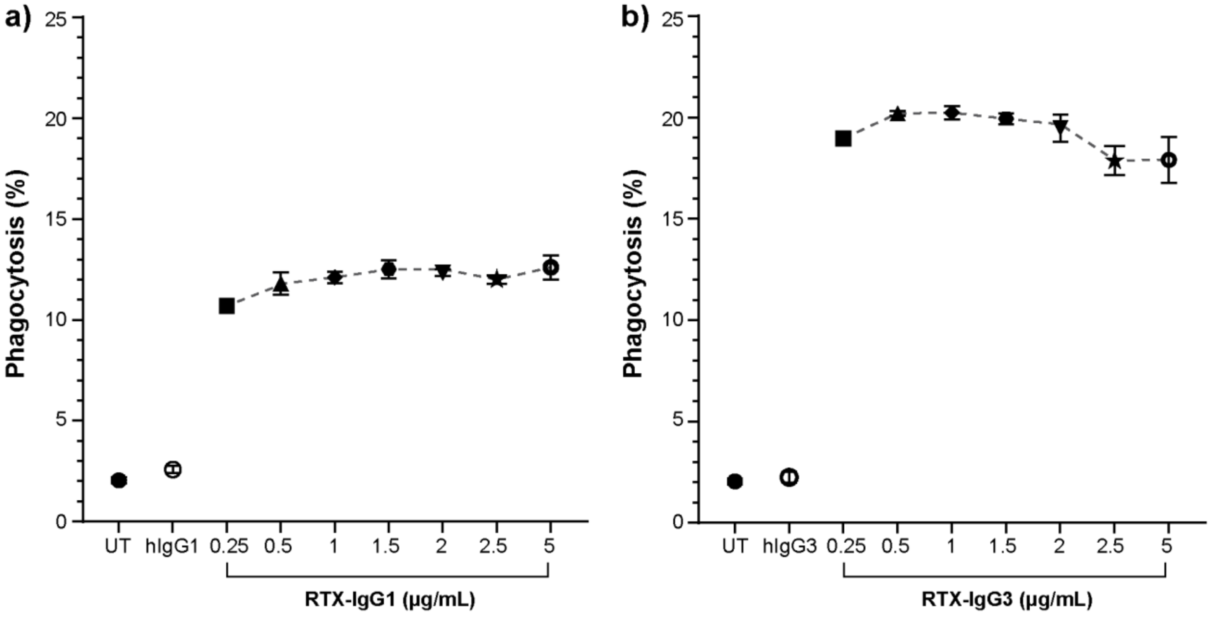


**Supplementary Fig. S4.** ADP of Granta-519 target cells, untreated (UT) or treated with RTX isotype or isotype control Ab, by MonoMac-6 effector cells (E:T ratio = 1:1). Percentage phagocytosis induced by different concentrations of a) RTX-IgG1 or b) RTX-IgG3. Results presented are mean ± SEM of three biological replicates.

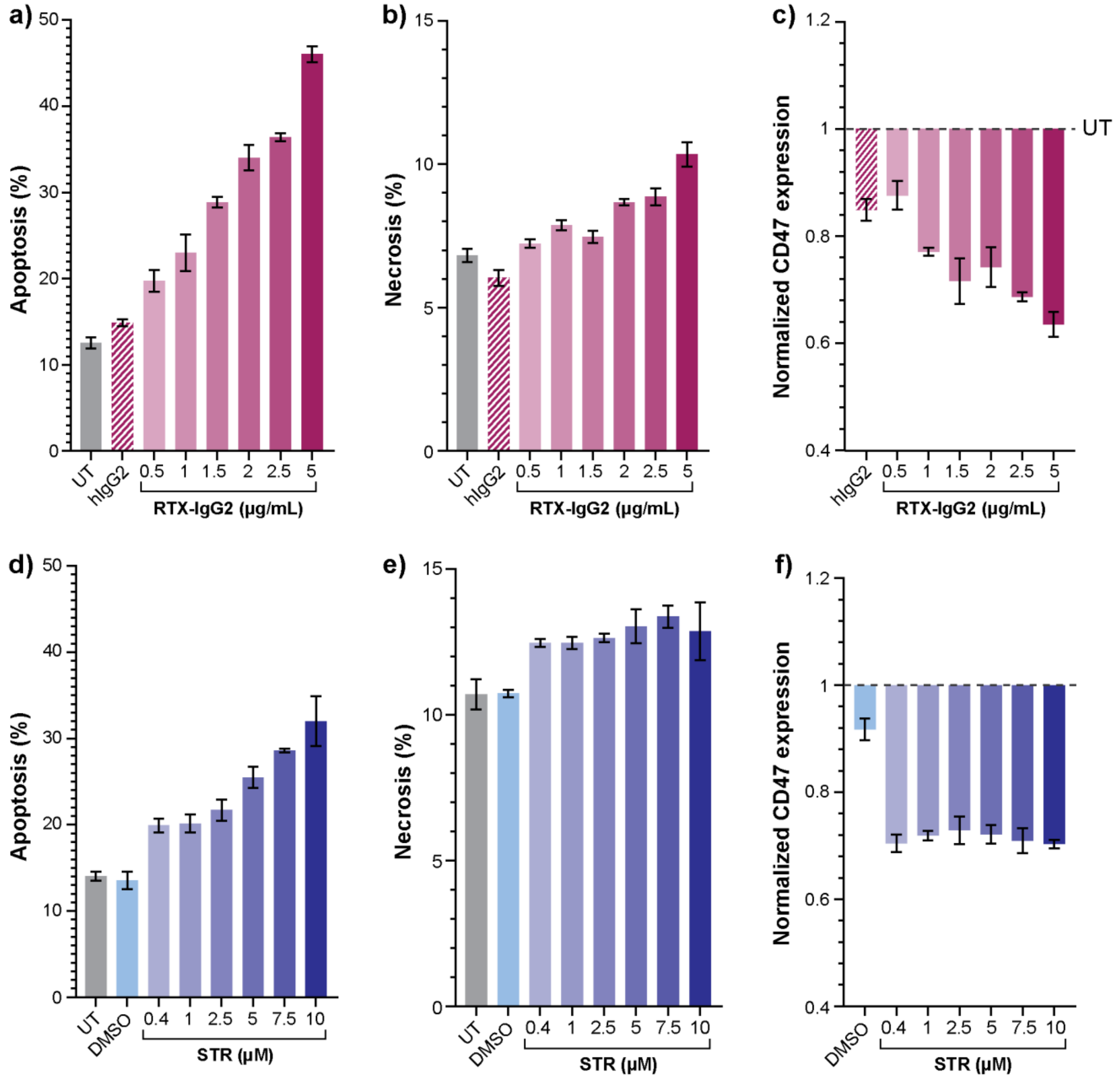


**Supplementary Fig. S5.** Analysis of apoptosis, necrosis and CD47 expression in Granta-519 cells treated with different concentration of a-c) RTX-IgG2 for 30 min or d-f) STR for 6 h before analysis. Data are presented as mean ± SEM of three biological replicates. Untreated cells (UT) and hIgG2 isotype control Ab were used as negative controls, while DMSO was used as vehicle control of STR treatment.

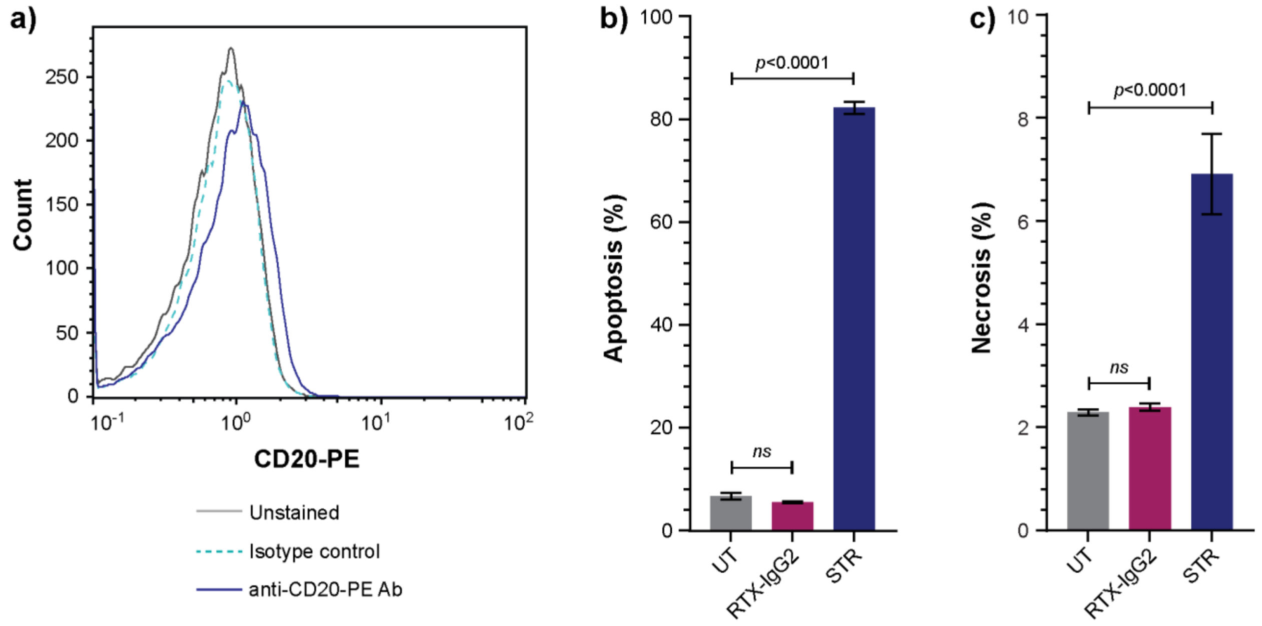


**Supplementary Fig. S6.** a) Representative histogram of CD20 staining of Reh cells. Analysis of b) apoptosis and c) necrosis in untreated (UT), RTX-IgG2 or STR-treated CD20-negative B-cells (Reh). Data are presented as mean ± SEM of three biological replicates. Statistical analysis by one-way ANOVA with Tukey-Kramer post-hoc test (ns = not significant).

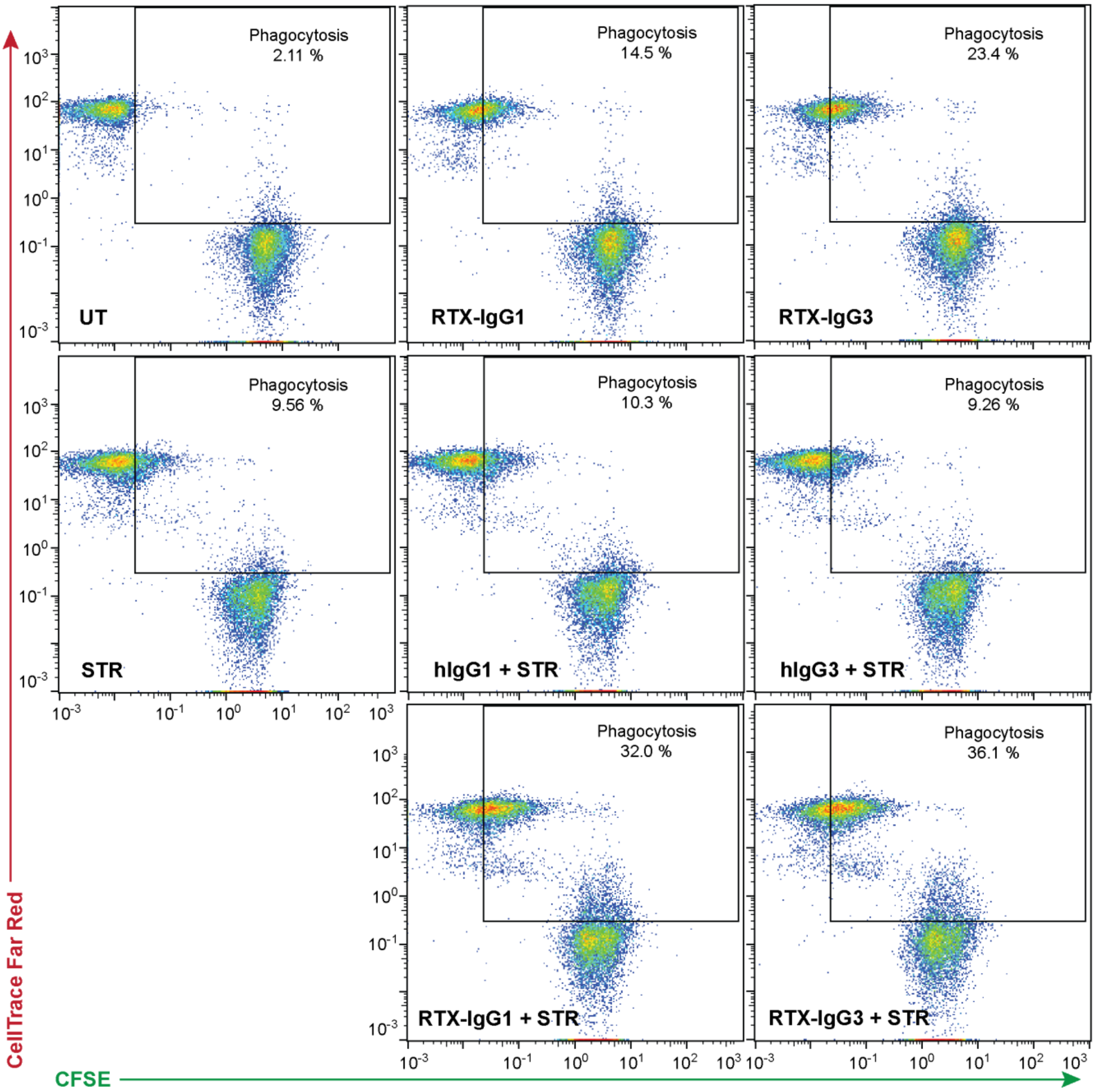


**Supplementary Fig. S7.** Representative flow plots showing ADP of Granta-519 cells treated with RTX-IgG1 or RTX-IgG3, and in combination with STR by MonoMac-6 cells (E:T ratio = 1:1). Untreated cells (UT) and STR in combinations with human Ab isotypes hIgG1 and hIgG3 were used as controls.

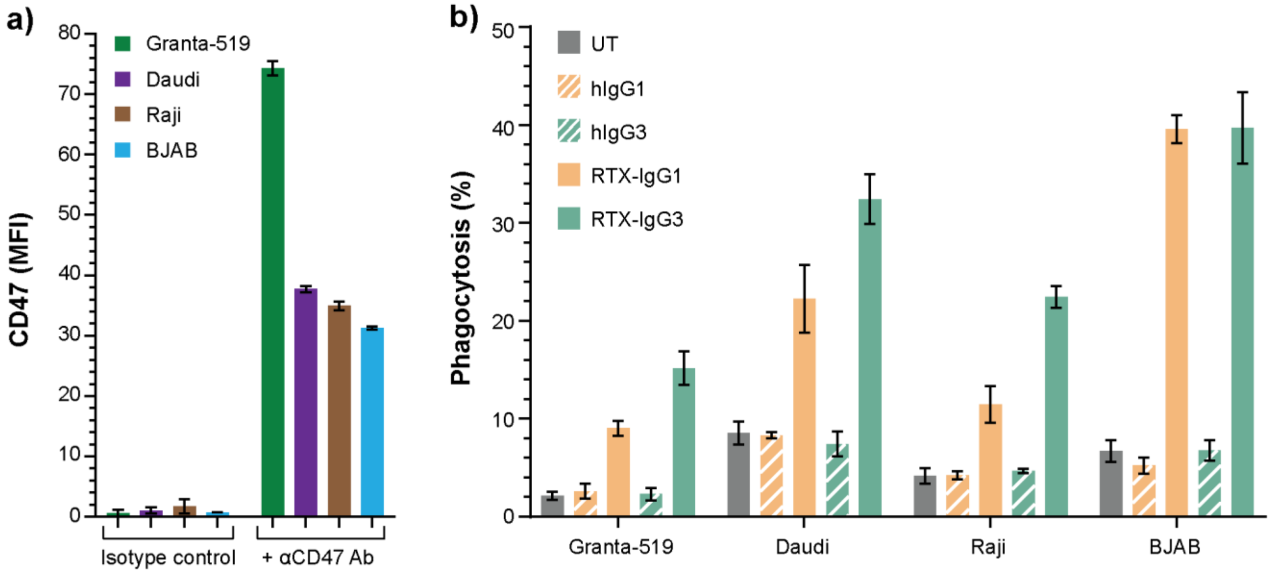


**Supplementary Fig. S8.** a) CD47 expression in four CD20$^{+}$ B-cell lymphoma cell lines. b) RTX-IgG1 and RTX-IgG3 mediated ADP in different B-cell lymphoma cell lines by MonoMac-6 effector cells (E:T ratio = 1:1). Data are presented as mean ± SEM of three biological replicates.

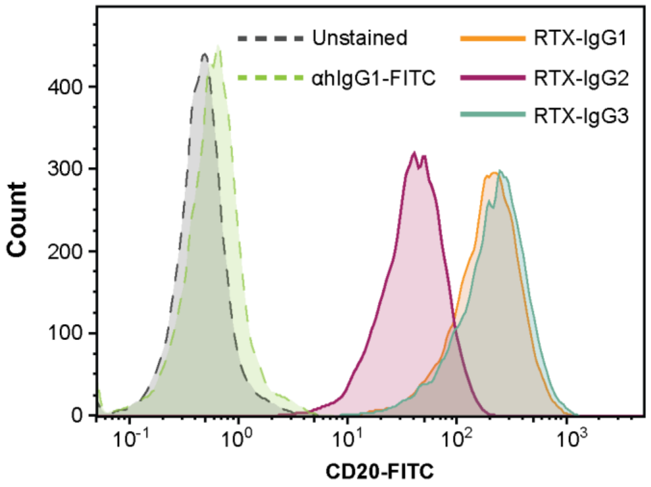


**Supplementary Fig. S9.** Binding of RTX-IgG1, RTX-IgG2 and RTX-IgG3 to CD20 on Granta-519 cells. Cells were incubated with RTX for 30 min at 37°C and subsequently stained with FITC-conjugated anti-human IgG Ab (αhIgG1-FITC).

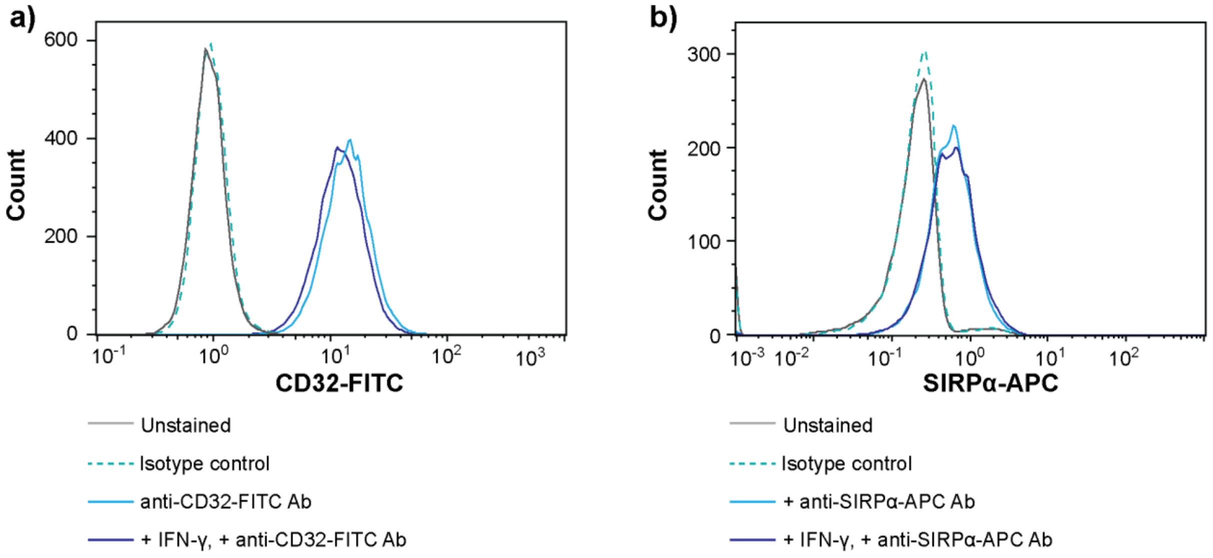


**Supplementary Fig. S10.** Representative histograms of a) CD32-FITC and b) SIRPα-APC staining of unstimulated or IFNγ-stimulated MonoMac-6 cells.

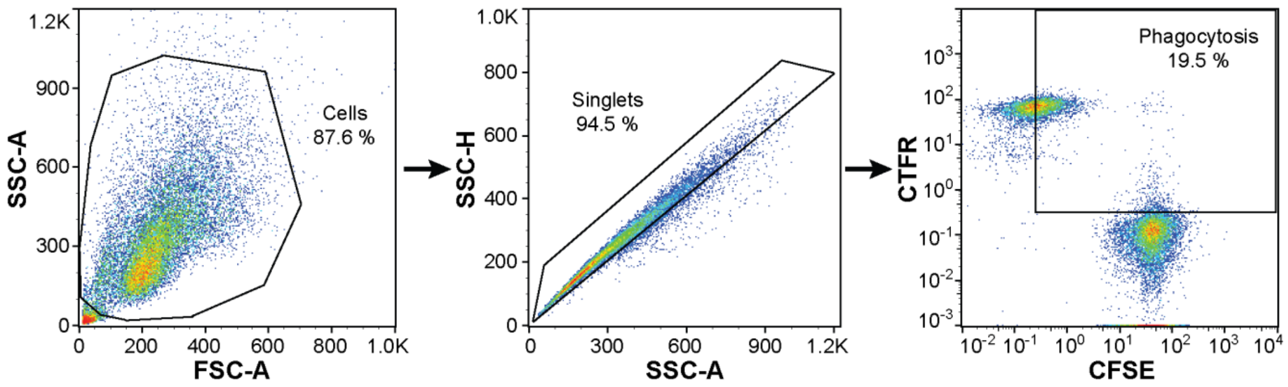


**Supplementary Fig. S11.** Gating strategy for quantitating phagocytosis. Phagocytosis was quantified as the percentage of double positive CFSE$^{+}$CTFR$^{+}$ MonoMac-6 cells (square gate). The gating was set based on unstained and single-colored stained controls. Results from one representative experiment out of three are shown.

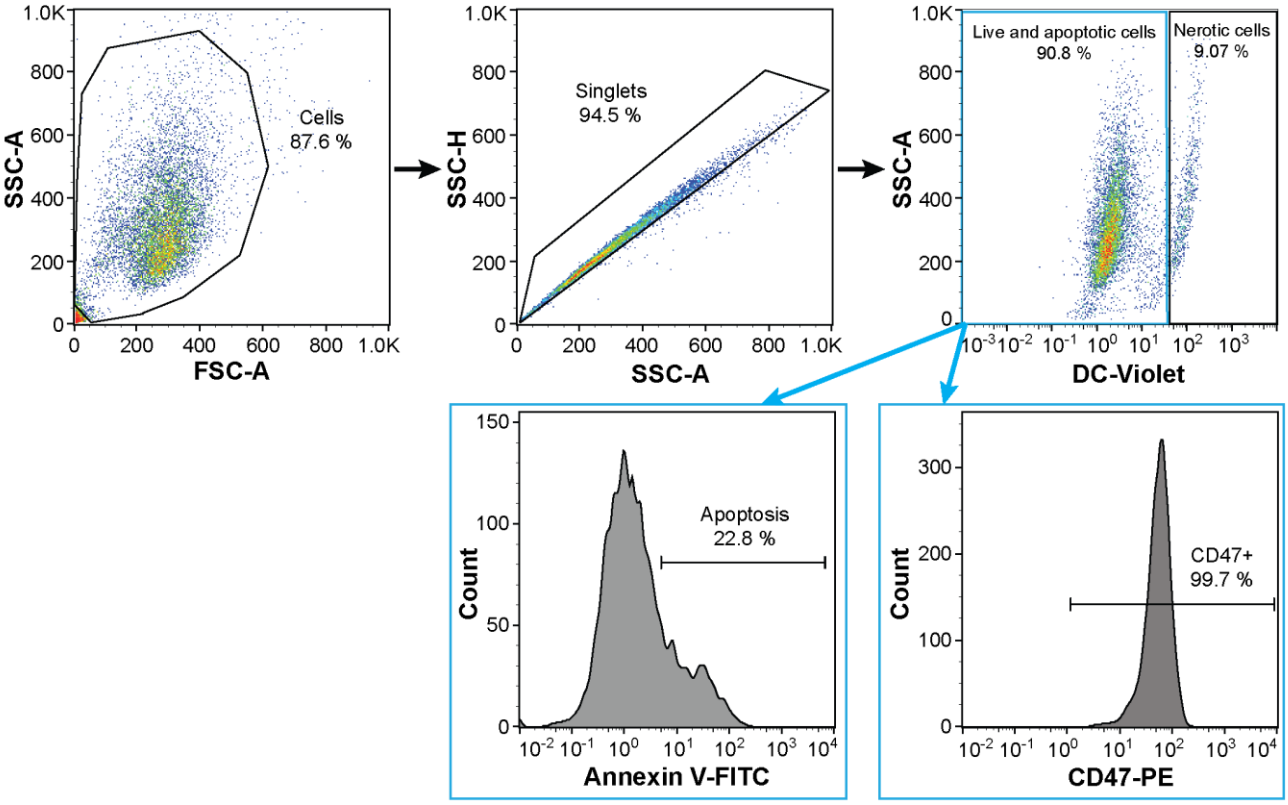


**Supplementary Fig. S12.** Gating strategy for apoptosis assay. Apoptotic cells were identified as Annexin V$^+$ DC-violet$^-$ cells, while double positive (Annexin V$^+$DC-violet$^+$) cells were regarded as necrotic cells with compromised cell membrane. Live and apoptotic cell population was further gated for CD47 expression using CD47-PE staining.
